# Bayesian inference of clonal expansions in a dated phylogeny

**DOI:** 10.1101/2021.07.01.450370

**Authors:** David Helekal, Alice Ledda, Erik Volz, David Wyllie, Xavier Didelot

## Abstract

Microbial population genetics models often assume that all lineages are constrained by the same population size dynamics over time. However, many neutral and selective events can invalidate this assumption, and can contribute to the clonal expansion of a specific lineage relative to the rest of the population. Such differential phylodynamic properties between lineages result in asymmetries and imbalances in phylogenetic trees that are sometimes described informally but which are difficult to analyse formally. To this end, we developed a model of how clonal expansions occur and affect the branching patterns of a phylogeny. We show how the parameters of this model can be inferred from a given dated phylogeny using Bayesian statistics, which allows us to assess the probability that one or more clonal expansion events occurred. For each putative clonal expansion event we estimate their date of emergence and subsequent phylodynamic trajectories, including their long-term evolutionary potential which is important to determine how much effort should be placed on specific control measures. We demonstrate the applicability of our methodology on simulated and real datasets.

## INTRODUCTION

In a microbial population, a clonal expansion event happens when a single individual (or clone) acquires an advantage relative to the rest of the population. This advantage could be selective, for example a mutation conferring antimicrobial resistance (Blair et al. 2015; Holmes et al. 2016), or neutral, for example a founder effect when the clone reaches a new population of susceptible hosts (Peter and Slatkin 2015). Whatever the mechanism, clonal expansion causes a single lineage to grow suddenly, leading to what were described as “epidemic clones” based on bacterial genotyping data (Maynard-Smith et al. 1993; Smith et al. 2003; Feil et al. 2004; Fraser et al. 2005). Since the advent of whole genome sequencing, clonal expansions have often been observed and described informally in pathogen phylogenetic trees, when a branch suddenly seems to split into multiple branches (McVicker et al. 2014; Holden et al. 2013; Eldholm et al. 2015; Shapiro 2016; Stoesser et al. 2016; Ledda et al. 2017).

Phylodynamics can be used to infer past population size changes given pathogen genetic data (Ho and Shapiro 2011; Volz et al. 2013). However, most phylodynamic methods assume that the same population size function applies to the whole population, which is inappropriate if a clonal expansion event affected only a subset of the sampled population. Differences between the branching observed in a phylogeny and the branching expected in the absence of any population structure can be used to test this assumption (Dearlove and Frost 2015; Volz et al. 2020). This principle provides a non-parametric approach to the detection of hidden population structure, based on rejection of the null hypothesis of an unstructured population. By contrast, here we develop and apply an explicit phylodynamic model for how structure arises through one or more clonal expansion events.

We describe a phylogenetic model of clonal expansion which is an extension of the coalescent framework (Kingman 1982; Donnelly and Tavare 1995; Rosenberg and Nordborg 2002), and more specifically an extension of the dated coalescent with heterochronous sampling and varying effective population size (Griffiths and Tavare 1994; Donnelly and Tavare 1995; Drummond et al. 2002, 2003; Biek et al. 2015). In brief, our population model consists of several subpopulations, including a “background” component of constant size, plus an unknown number of additional components each of which corresponds to a clonal expansion event, with an associated time of emergence, growth rate and maximum population size (carrying capacity). We also describe how to perform Bayesian inference under this model, taking as input a dated phylogeny, such that can be reconstructed using BEAST (Suchard et al. 2018), BEAST2 (Bouckaert et al. 2019), treedater (Volz and Frost 2017), TreeTime (Sagulenko et al. 2018) or BactDating (Didelot et al. 2018). In this inferential setting, our methodology allows us to detect putative clonal expansions, assess their statistical significance and the specific parameters controlling their growth. We performed inference on simulated datasets, where the correct clonal expansions that took place are known, in order to benchmark the specificity and sensitivity of our methodology. We also analysed several real datasets from recent studies on infectious diseases, and show that our new method can reveal important features in pathogen evolutionary epidemiology that would otherwise be difficult to analyse.

## MATERIALS AND METHODS

### Mathematical model description

We consider the ancestry of a sample of *N* individuals indexed by *i* ∈ {1,…,*N*}, with sampling times denoted **t** = {*t_i_*}_*i*∈{1,…,*N*}_. Here and elsewhere in this article, time is measured backward in time so that for example if *t*_1_ < *t*_2_ then sample 1 is more recent than sample 2. The population is structured into *M* ≥ 1 subpopulations indexed by *j* ∈ {1,…,*M*}: the subpopulations *j* ∈ {1,…,*M*−1} correspond to *M*−1 “clonal expansion” subpopulations whereas the population *j* = *M* is called the “background” subpopulation. Each individual has the same probability *θ_j_* of belonging to subpopulation *j*, with ***θ*** = {*θ*_1_,…, *θ_M_*} and 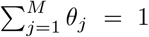. This population structure therefore partitions the sampled individuals {1,…,*N*} into *M* mutually disjoint subsets **f** = {*f*_1_,… *f*_*M*−1_, *f_M_*} with 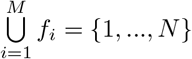.

The background subpopulation (*j* = *M*) is assumed to be ruled by the coalescent process with constant population size *N_M_* (Kingman 1982). Each of the other subpopulations (*j* = 1,…, *M* − 1) on the other hand is ruled by a coalescent model with its own varying population size function (Griffiths and Tavare 1994). For each of these clonal expansion subpopulations we define a time of emergence 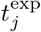, a carrying capacity *N_j_* and the time *h_j_* it takes to reach half of the carrying capacity. Together these parameters determine the size *α_j_*(*t*) of the subpopulation *j* at time *t* as follows:

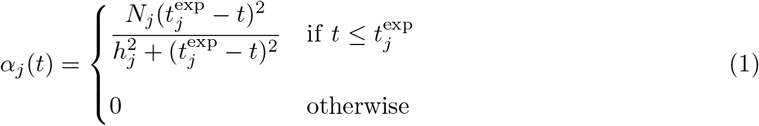

Note that this function has the property 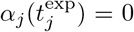 so that the population size reaches zero, when the expansion begins at 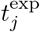. This forces the coalescent rate for a lineage to diverge to infinity as 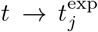. As such all lineages from the subpopulation are forced to coalesce before 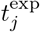. From a modelling perspective this can be interpreted as the population being negligible at the time of the lineage diverging. Furthermore, *α_j_*(*t*) → *N_j_* when *t* → −∞ in accordance with the definition of a carrying capacity being the size reached in the long term. Finally we note that 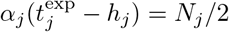, which means that hj is indeed the time it takes to reach half of the carrying capacity. This function represents a qualitative approximation to the population dynamics of a clonal expansion.

To complete the definition of the joint ancestral process for all *N* individuals, we consider that each of the clonal expansions originated from either the background subpopulation or from one of the preexisting clonal expansions. Let *d_j_* denote the population from within which expansion *j* ∈ {1,…, *M* − 1} originates. Therefore *d_j_* ∈ {1,…,*M*} with the condition that if *d_j_* < *M* then 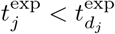 (if the origin is not the background subpopulation, it is another clonal expansion that must have emerged beforehand). Since each expansion starts with a negligible population size, this implies that the group of leaves sampled from a subpopulation is either monophyletic (if this subpopulation is not the origin of another one) or paraphyletic (otherwise) in the phylogeny of all *N* individuals.

Table 1 summarises the parameters involved in this model, and lists the priors which were used to perform Bayesian inference under this model. The background population size effectively acts as a scale parameter on the entire process. First of all, we assume that the final effective population sizes of the individual expansions are in the same order of magnitude as the background population size, as defined by the prior probability *π*(*N_j_* | *N_M_*). Furthermore, by affecting the expected time to most recent ancestor of the phylogeny, the background population size strongly determines which clonal expansions will be detectable and which will not. An expansion which occurred in the distant past, or whose growth rate is slow is very likely to fully coalesce while its effective population size remains near constant, making it undetectable. As such we condition both 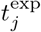 and *h_j_* on *N_M_*, leading to the prior distributions 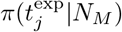 and *π*(*h_j_*|*N_M_*).

**Table 1:**
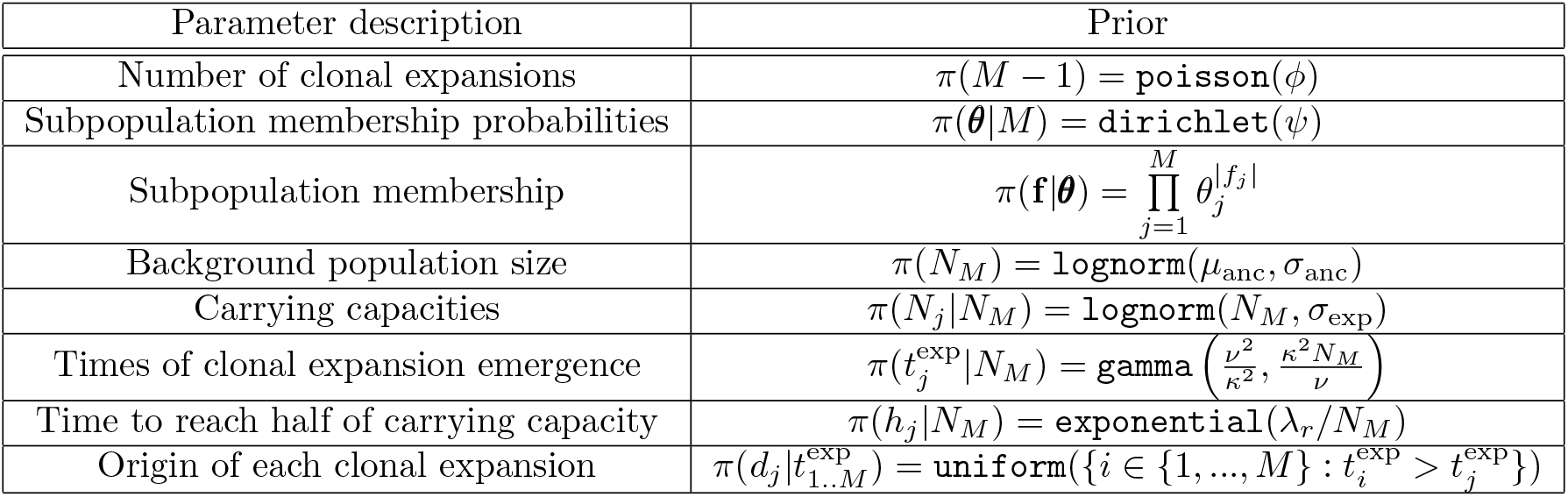
Summary of parameters and priors used for Bayesian inference

### Bayesian inference

Performing inference under the clonal expansion model above for a given dated phylogeny **g** requires estimation of the value of all the underlying parameters of this model, including the unknown number of subpopulations *M*. We consider the prior distributions summarised in Table 1. For convenience, let ***α*** denote the combination of the parameters *N_M_* for the background population and 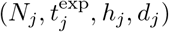 for each of the *j* = 1,…, *M* − 1 clonal expansions. The joint prior on ***α*** is therefore:

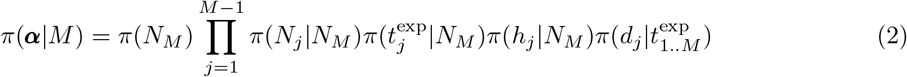

We can decompose the posterior probability of the model parameters given the dated phylogeny as follows:

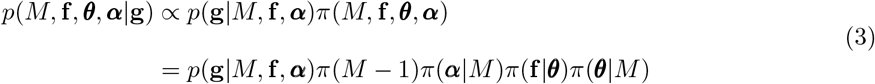

All other terms correspond to prior densities given in Table 1 and Equation 2, except for the first term *p*(**g**|*M*, **f**, ***α***) which is the likelihood of the dated phylogeny when all parameters are known, including which leaves belong to which subpopulations, the population size function of each subpopulation, and the origin of each clonal expansion subpopulation. In these conditions the likelihood is simply the product of likelihoods of the coalescent process in each of the subpopulations. Note that as *M* increases, meaning that more clonal expansion events are introduced, the probability *π*(**f**|***θ***) decreases since the number of possible membership assignment increases, but this is compensated by an increase in the likelihood *p*(*M*, **f**, ***θ, α***|**g**) since coalescent events between lineages in different components become disallowed. Let **g**_*j*_ denote the part of the dated phylogeny that corresponds to the subpopulation *j*. Knowledge of (*M*, **f**, ***α***) allows us to decompose exactly the genealogy **g** into each of the **g**_*j*_ components. Note in particular that a component **g**_*j*_ contains all the leaves indexed in *f_j_* plus a leaf dated at 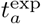 for each subpopulation *a* such that *d_a_* = *j*, meaning that the origin of *a* is *j*. With these notations, the likelihood is therefore decomposed as:

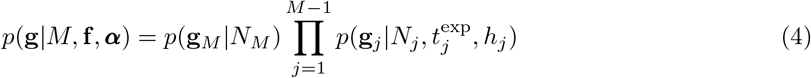

The first term corresponds to the coalescent process in the background subpopulation, with constant population size *α_M_*(*t*) = *N_M_*, and the remaining terms correspond to the coalescent process in the clonal expansion subpopulations, each with their own population size function *α_j_*(*t*) as defined in Equation 1. These terms can be computed using standard coalescent theory (Griffiths and Tavare 1994; Donnelly and Tavare 1995; Drummond et al. 2002). Briefly, if a population has size *α*(*t*) and *A*(*t*) extent lineages at time *t*, then the probability of a dated phylogeny **g** with *n* − 1 coalescent events at times *c*_1_,…, *c*_*n*−1_ is given by:

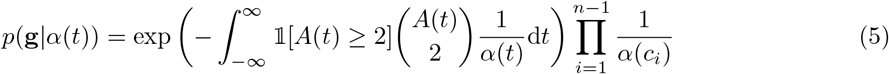

Note the absence of the 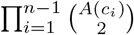 term as this is the likelihood of the entire genealogy, meaning both the branch lengths and the topology, so that this term from the probability of the waiting times cancel out with its reciprocal from the probability of the topology.

The computation in Equation 5 requires us to calculate the integral of the reciprocal of the population size function, for each interval of time in which *A*(*t*) is constant and greater than one. This is straightforward for the background subpopulation, and for each clonal expansion subpopulation *j* with the population size function given in Equation 1 we can use the primitive function:

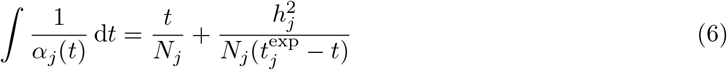

This completes the definition of the posterior probability in Equation 3. In order to sample from this posterior distribution, we use a Reversible jump Markov Chain Monte-Carlo (Green 1995; Hastie and Green 2012), since the dimensionality of the parameter space depends on the unknown parameter *M*. The details of the updates used in this procedure are given in Supplementary Material. Unless otherwise stated, during inference on all real and simulated datasets, we used the following hyperparameters: *θ* = 1 *ϕ* = 1 *μ*_anc_ = 3 *σ*_anc_ = 3, *σ*_exp_ = 1, *ν* = 1/2, *κ* = 1/2, λ_*r*_ = 5.

### Practical considerations

Here we provide a practical summary of the model hyperparameters, with advice on how to elicit them, as well as considerations for the input phylogeny. The parameter *ϕ* corresponds to the Poisson mean of the prior placed on the number of expansions. The parameter *ψ* corresponds to the concentration parameter of a Dirichlet prior on subpopulation membership probabilities, and therefore regulates how balanced the number of tips assigned to individual subpopulations will be. *μ*_anc_ and *σ*_anc_ correspond to the log-normal mean and standard deviation of the prior placed on the effective population size *N*_M_ of the background population. The prior distribution on the expansion parameters are set so that *N*_M_ acts as a scale parameter for the entire process. *σ*_exp_ corresponds to the log-normal standard deviation for the effective population size of expansions, with their log-normal mean being *N_M_*. The parameters *ν* and *κ* determine the mean *νN_M_* and variance (*κN_M_*)^2^ of the expansion emergence time prior. Finally, the parameter λ_*r*_ controls the mean *N_M_*/λ_*r*_ of the prior distribution on the time it takes an expansion to reach half of its carrying capacity.

An important practical aspect of Bayesian inference is elicitation of priors. While we provide a set of default values which should be a reasonable starting point for most applications, we encourage to consider the specificities of each application. The default hyperparameter values are *ϕ* = 1, *ψ* = 2, *μ*_anc_ = 3, *σ*_anc_ = 3, *σ*_exp_ = 1/2, *ν* = 1/2, *κ* = 1/2, λ_*r*_ = 5. When considering a specific application, particular attention should be given to hyperparameters *ϕ* and *σ*_exp_. We advise starting with *ϕ* = 1, and adjusting upwards if there is a reasonably strong belief that the phylogeny may contain a large number of clonal expansions, for example if the samples are clustered across several geographically disconnected locations, as these processes are likely to give rise to clonal expansions. To elicit *σ*_exp_, one should consider what the effective population size of clonal expansions can be relative to the size of background population. In general *σ*_exp_ < 1 to penalise unreasonably large carrying capacities which could lead to identifiability issues. Finally *μ*_anc_ and *σ*_anc_ can be adjusted to be make the prior on *N_M_* more informative if we have prior knowledge on the background population size. The concentration parameter *ψ* can be adjusted upwards to discourage expansion that consist of only a few tips.

Our model assumes that the input phylogeny is correct, and inaccuracies will affect the inferences on clonal expansions. In particular it is important to pay attention to unrealistic branch lengths. Negative branch lengths outright invalidate our approach as they are not consistent with the coalescent framework. With maximum likelihood trees, unrealistically short or zero branch lengths could lead to false identification expansions. When using a Bayesian phylogenetic reconstruction, care should be taken that the summarised phylogeny has all branch lengths strictly positive. The computational time required per iteration scales linearly with the number of tips in a phylogeny. However, mixing properties and number of iterations required to reach satisfactory results generally depend on the complexity of the underlying population structure, as well as the compatibility of the phylogeny with our model. Posterior distributions under the model are relatively complex and high-dimensional, which makes their analysis a non-trivial task. The posterior probability that a pair of tips belongs to the same population partition block can be evaluated and visualised as a heat map whose block structure coincides with the posterior clonal expansion structure, while also including information about the underlying uncertainties. Combined with information from the posterior marginal for the number of clonal expansions, different expansion scenarios can then be formulated and evaluated.

### Simulation of testing data

The process characterised above represents a standard Continuous Time Markov Chain (CTMC) and as such can be simulated directly via Gillespie’s algorithm (Gillespie 1976). The waiting times are sampled through inverse transform sampling with the inverse of the total process rate being approximated numerically. For the simulation of the genealogy in the first illustrative dataset presented, we used the following hyperparameters: *θ* = 1, *ϕ* = 2, *μ*_anc_ = 4, *σ*_anc_ = 1/2, *σ*_exp_ = 1, *ν* = 1/2, *κ* = 1/4, λ_*r*_ = 5. For all other simulated genealogies we used: *θ* = 1, *ϕ* = 2, *μ*_anc_ = 5, *σ*_anc_ = 1/2, *σ*_exp_ = 1/2, *ν* = 1/3, *κ* = 1/4, λ_*r*_ = 5.

### Implementation

We implemented the simulation and inference methods described in this paper into a new R package entitled *CaveDive* which is available at https://github.com/dhelekal/CaveDive. The package uses ape (Paradis and Schliep 2019) as a backend for handling phylogenies and ggtree (Yu et al. 2017) for handling the visualisation of results. We also used the coda package (Plummer et al. 2006) to assess the convergence and mixing properties of our MCMC sampler, and found them to be satisfactory with Gelman-Rubin statistics being less than 1.1 and the effective sample sizes in excess of 200 for all parameters in the runs presented below. All runs were performed on a single core of Intel(R) Core(TM) i7-3770 CPU with 8GB RAM.

## RESULTS

### Illustration of the clonal expansion model

In order to illustrate the concepts behind our clonal expansion model, we simulated from it the scenario shown in Figure 1. In this example the population was made of *M* = 4 components: a background subpopulation (pink) and three clonal expansions (blue, orange, green). Figure 1A shows the effective population size of the four subpopulations as a function of time. The background subpopulation remains of a constant size throughout, whereas each of the clonal expansions is characterised by a time when the expansion started, a carrying capacity and a time to reach half of this carrying capacity. The blue clonal expansion was the first one to have emerged, it has a large carrying capacity but this potential is almost fully realised. The orange clonal expansion emerged next and very quickly reached a relatively small carrying capacity. Finally, the green clonal expansion emerged and at the present time it is still growing and far from having reached its capacity.

**Figure 1:**
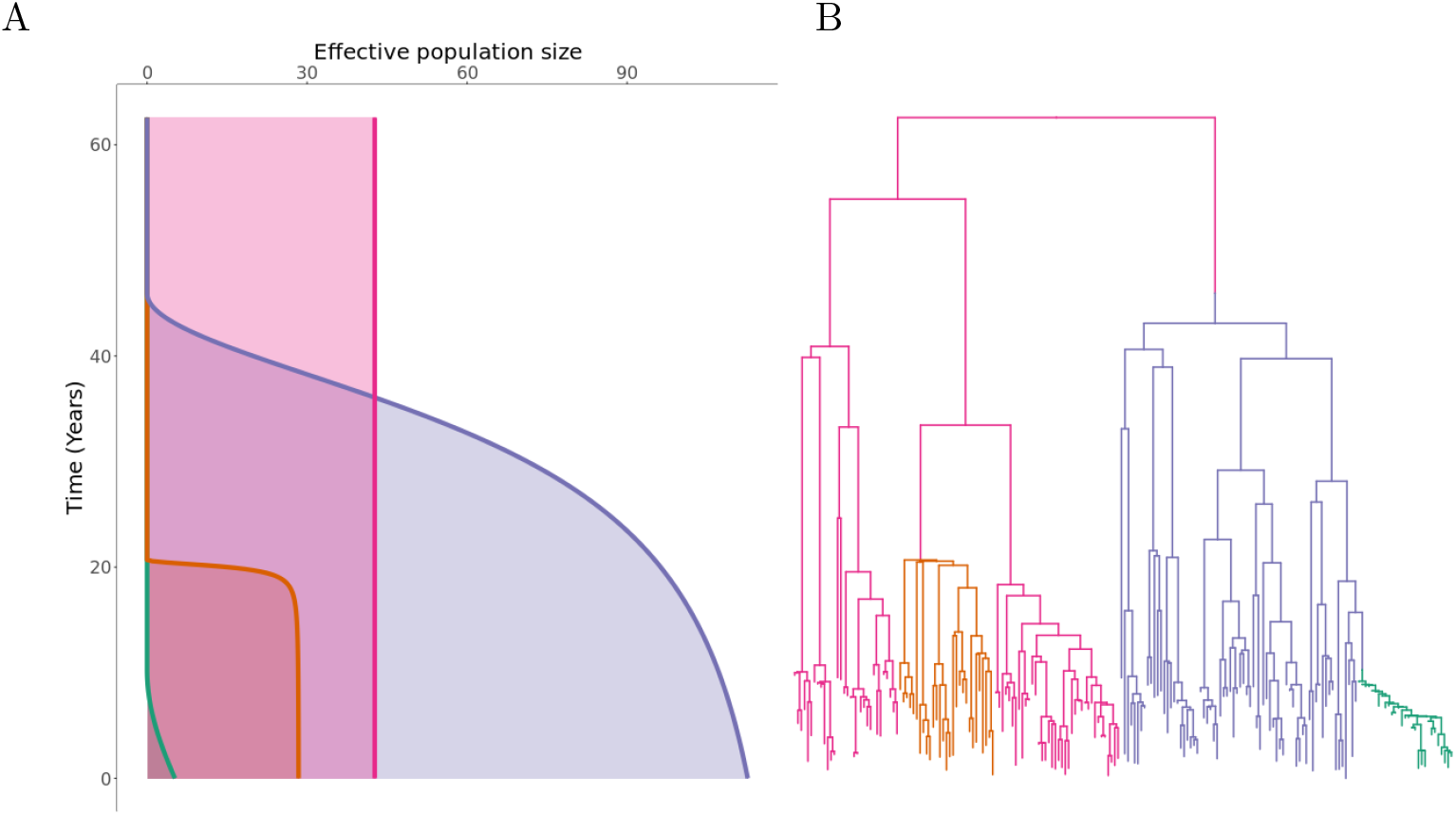
A realisation from the clonal expansion model. (A) Population size functions for each of the subpopulations. (B) Dated phylogeny coloured according to subpopulation as in part (A).

Figure 1B shows the corresponding dated phylogeny with 200 tips that was simulated in this example. Each point on this dated phylogeny belongs to one of the subpopulations and is coloured accordingly as in Figure 1A. A change of colour therefore corresponds to the emergence of a clonal expansion. The blue and orange clonal expansions emerged out of the background subpopulation, whereas the green expansion emerged out of the preexisting blue expansion, as can be seen from the transition from blue to green.

For each of the four subpopulations, the population size function (Figure 1A) determines the branching pattern in the corresponding part of the phylogeny (Figure 1B). For example, the background subpopulation (pink) had a constant population size and the corresponding branches are therefore consistent with expectation under the standard coalescent model. By contrast, the three clonal expansions have been growing in size more or less suddenly resulting in star-like branchings soon after their times of emergence. The orange and blue clonal expansions have almost reached their carrying capacities so that recent branchings are similar to the expectation under a constant population size as for the background subpopulation. The green clonal expansion on the other hand is still growing and remains very small giving it a more linear structure.

### Application to a single simulated dataset

We attempted to reconstruct the clonal expansion structure underlying the example shown in Figure 1. In this inferential setting, the input data is therefore the dated phylogeny shown in Figure 1B, without the colouring or location of colour changes that correspond to the emergence of clonal expansions. The aim is to infer the correct number of clonal expansions (three in this case), their locations on the phylogeny (colour changes in Figure 1B) as well as the demographic properties of each subpopulation (Figure 1A).

The priors used during the inference were the same as used for the simulation of this phylogeny. The MCMC sampler was run for 10^7^ iterations with sampling every 1000 iterations, which took approximately 1.5 hours. The results are shown in Figures 2 and S1. The correct number of three clonal expansions was inferred with 67.5% of the posterior probability mass concentrated there, and the majority of the remainder of the posterior probability mass shared between four and five clonal expansions (Figure 2B). This suggests that although the phylogenetic data is informative about the three correct expansions, it is not possible to rule out the existence of other expansions that would have left little effect on the phylogeny, for example if they were very recent and if they would have concerned only a small number of leaves. The correct position for the clonal expansions was inferred with high probability, although it was not always possible to distinguish with certainty between the correct branch or the ones directly above or below (Figure 2C-D). The demographic parameters of the three clonal expansions (carrying capacity and time to reach half of it) were also correctly inferred, resulting in posterior distributions for the effective population size of each expansion over time similar to the ones used in the simulation (Figure 2E-G). The only exception concerned the carrying capacity parameter of the orange expansion which was slightly overestimated (branch 49, cf Figure 2F), because of the difficulty in correctly inferring such a sudden and self-limiting expansion. For comparison purposes, we applied treestructure (Volz et al. 2020) to the same dataset, and found that only the most recent clonal expansion was detected (Figure S2). We also applied treeImbalance (Dearlove and Frost 2015) which found several nodes with statistically significant evidence of imbalance (Figure S3).

**Figure 2:**
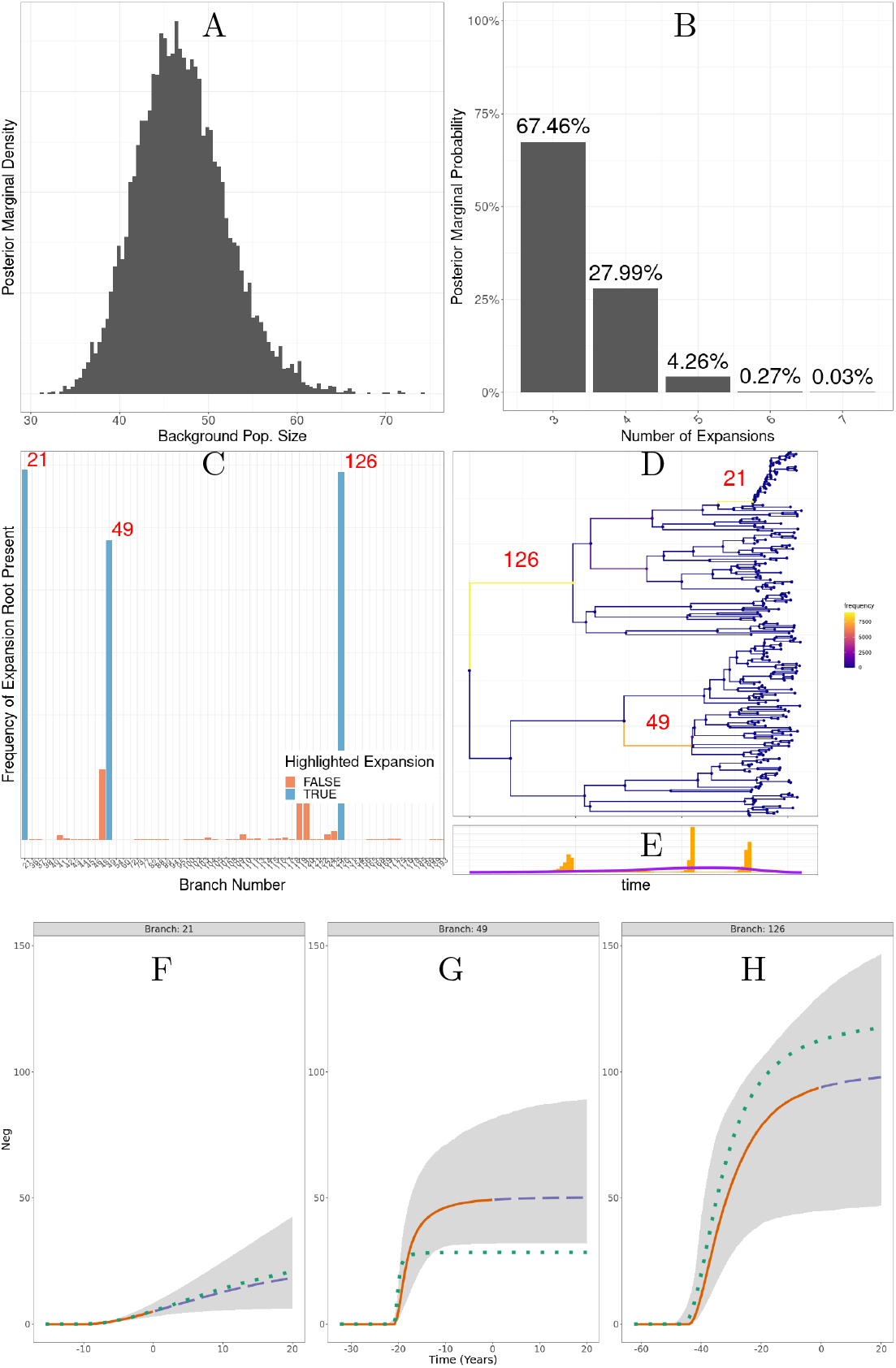
Application to the simulated dataset shown in Figure 1. (A) Posterior distribution of the background population size. (B) Posterior distribution of the number of clonal expansions. (C-D) Posterior probabilities of having a clonal expansions on different branches of the tree, with the indexes of three branches of interest shown. (E) Posterior distribution of clonal expansion starting times, with prior shown in purple. (F-H) Posterior reconstruction of the expansion population dynamics. 95% credible intervals in grey. Median in solid orange for past population dynamics and dashed blue for future prediction of the population dynamics. True population dynamics in dotted green.

### Application to multiple simulated datasets

Firstly we performed inference based on 100 simulated dated phylogenies, in which no clonal expansion event occurred, so that the whole phylogeny is ruled by a single coalescent process with constant population size. Each phylogeny had a number of tips uniformly sampled between 80 and 300. This allowed us to evaluate the false discovery rate of our methodology. For each dataset in this test, the MCMC was run for 10^6^ iterations with sampling every 100 iterations. We found that in 98% of the replicates, the highest posterior probability was of having no clonal expansion, corresponding to a 2% false positive rate. Such occasional false positive detection of clonal expansion events is to be expected due to the fact that such events can leave little phylogenetic signature, and therefore be difficult to rule out.

Secondly we performed inference based on 200 simulated dated phylogenies with 100 tips each, in which a single clonal expansion event occurred, and the results are shown in Figure 3. In this benchmark, the MCMC was run for 10^7^ iterations with sampling every 1000 iterations. For nearly 74.5% of the simulated datasets a single clonal expansion was found to be most likely (Figure 3A), as was indeed correct. In 15.5% of the replicates no clonal expansion was found to be most likely, indicating a false negative case. This result reflects the fact that some clonal expansion events are hard to infer if they left little phylogenetic signature, for example if they occurred very recently, were sampled only a small number of times, or occurred so long ago that almost all coalescent events occur before the period of rapid growth. Finally, in 10% of the simulated datasets two clonal expansions were found to be most likely, representing a relatively low rate of false positive detection, for the same reasons as in the previous simulations where no clonal expansion had happened.

**Figure 3:**
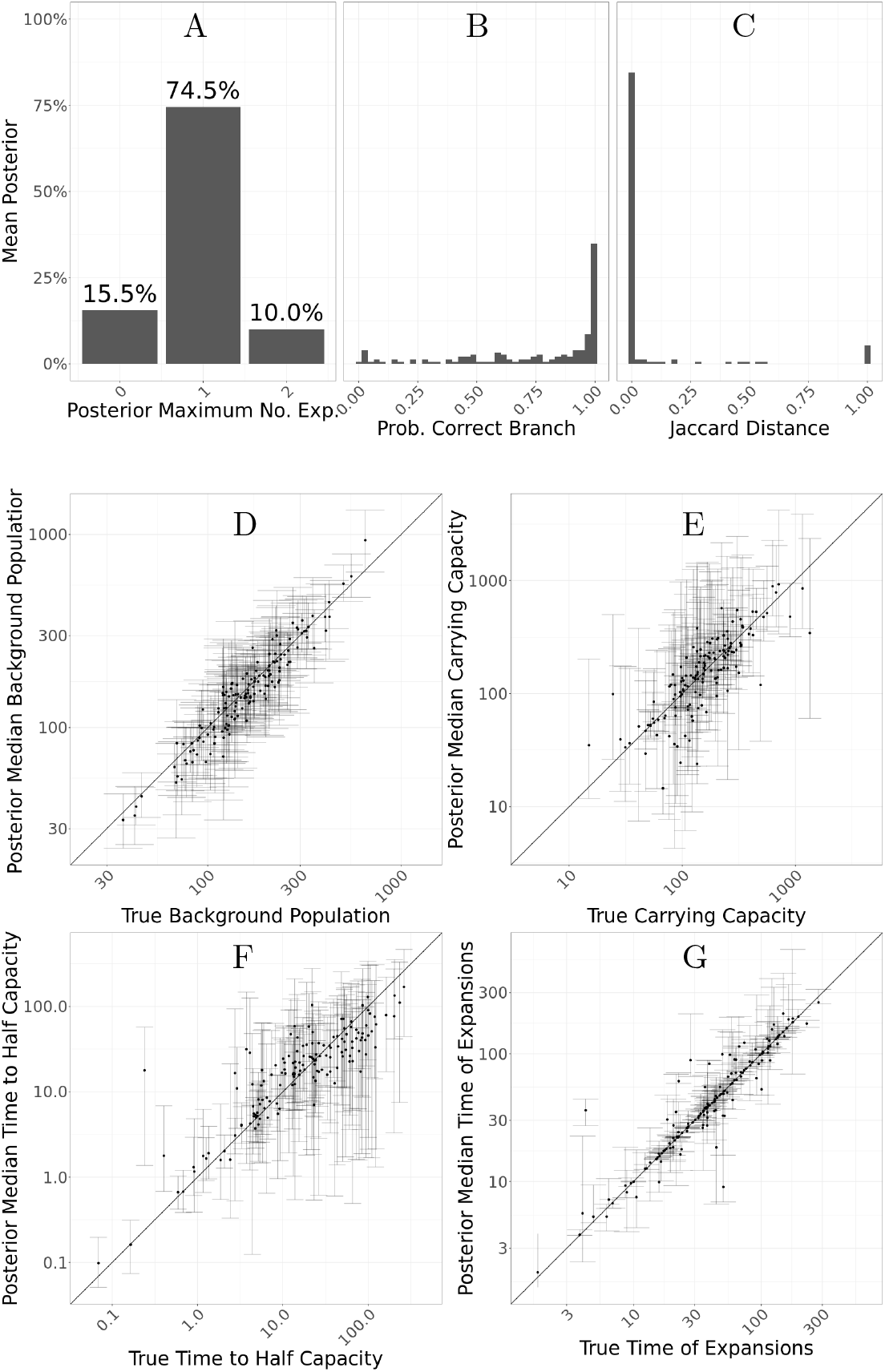
Application to 200 simulated trees containing one expansion. (A) Histogram of posterior modes for the number of expansions. (B) Histogram of probability to have a clonal expansion on the correct branch. (C) Histogram of Jaccard distances between the true expansion and the expansion corresponding to the mode branch. (D-G) Scatter plots showing posterior median and 95% credible interval for individual expansion parameters, with correct values on the x-axis and inferred values on the y-axis. Parts B-G only include simulations where the inferred mode of the number of expansions was one.

When a single clonal expansion was inferred, the probability of having this inferred event on the correct branch was typically high (Figure 3B). However, when that was not the case, the clonal expansion was almost always inferred on a very closely related branch, as can be seen when computing the Jaccard distance between the correct and inferred expansion memberships (Figure 3C). The inferred effective population size of the background population was highly consistent with the correct values (Figure 3D), and the same was true for the carrying capacity of the clonal expansion (Figure 3E). The time taken to reach half of the carrying capacity was harder to infer, with little correlation between the correct and inferred values (Figure 3F). The dating of the emergence of the clonal expansion was often very precisely estimated (Figure 3G), although in some cases the credible interval on this parameter was larger, which would be expected for example if the clonal expansion happened on a long branch.

Finally we performed inference based on 100 simulated dated phylogenies in which two or more clonal expansion events occurred. We have simulated four sets of 25 phylogenies, with each set having two, three, four, and five expansions respectively. These particular phylogenies were simulated using a total number of tips equal to 60 plus 40 times the number of expansions. In this benchmark, the MCMC was run for 2 × 10^7^ iterations with sampling every 2000 iterations. The expected posterior (Figure 4A) marginals for the number of expansions show a clear trend in probability mass being located on a greater number of putative clonal expansions as the number of simulated expansions increases. We observe a tendency to underestimate the number of expansions, which increases with the true number of expansions. In terms of the posterior expectation of the number of expansions (Figure 4B) we observe a clear increasing trend in terms of the medians, which initially closely follow the true number of expansions in the case of two and three expansion phylogenies, and underestimates the number of expansions for phylogenies with four and five expansions. This result reflects our relatively conservative prior on the number of expansions *M* ∝ Poisson(1), and the fact that they become harder to detect as more and more occur on the same phylogeny, frequently with some expansions originating from within another.

**Figure 4:**
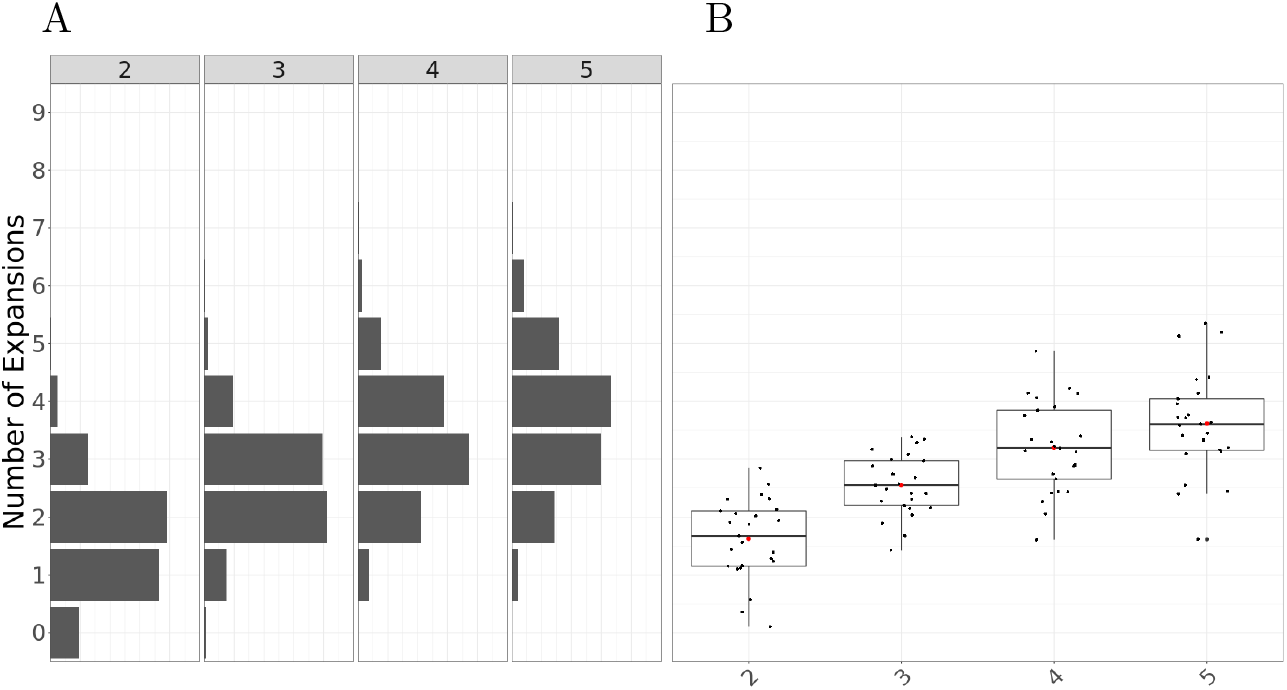
Application to 100 simulated datasets, with 25 per each scenario with 2, 3, 4 and 5 expansions. (A) Expected posterior distributions for the number of expansions for each scenario. (B) Box plots of the posterior mean number of expansions for each simulation by scenario.

### Application to *Streptococcus pneumoniae* dataset GPSC18

As the first real dataset to demonstrate our method, we used a global collection of genomes from the Global Pneumococcal Sequence Cluster 18 (GPSC18) from a previously published study (Gladstone et al. 2019). In this study, the authors described increased invasiveness in serotype 14 compared to the background genotypes in the GPSC18 cluster. Indeed, serotype 14 is one of the leading causes of invasive pneumococcal disease (Song et al. 2013), and its prevalence was reported to have increased in recent years, despite its inclusion in pneumococcal conjugate vaccines (He et al. 2015). This dataset consists of 228 genomes collected between 1991 and 2015, for which a dated phylogeny has been previously published (Gladstone et al. 2020). Running our software for 10^8^ iterations took approximately 15 hours. The results are shown in Figures 5 and S4. The posterior inferred under our model includes a single clonal expansion with very high certainty (Figure 5A), although other less certain expansions can not be completely ruled out. The model therefore separates the genomes into two categories, with about 80% of them belonging to the expansion and the remainder belonging to the background population (Figure 5B). Notably, the expansion contains the vast majority of serotype 14 isolates, while containing only very few isolates corresponding to other serotypes (Figure 5C). Conversely, the background population contained few isolates of serotype 14, with most of them being of serotype 7C, 16F, 19A or 19F (Figure 5C). The inferred population size dynamics of clonal expansion suggests that currently the expansion is of a slightly smaller size than the background population of the GPSC18 cluster, but that it it is still growing and might increase beyond the size of the background population in the future (Figure 5D). This result is consistent with the fact that more genomes belonged to the clonal expansion than to the background population: since serotype 14 is more associated with disease, it would tend to be overrepresented in isolate collections (Didelot and Maiden 2010).

**Figure 5:**
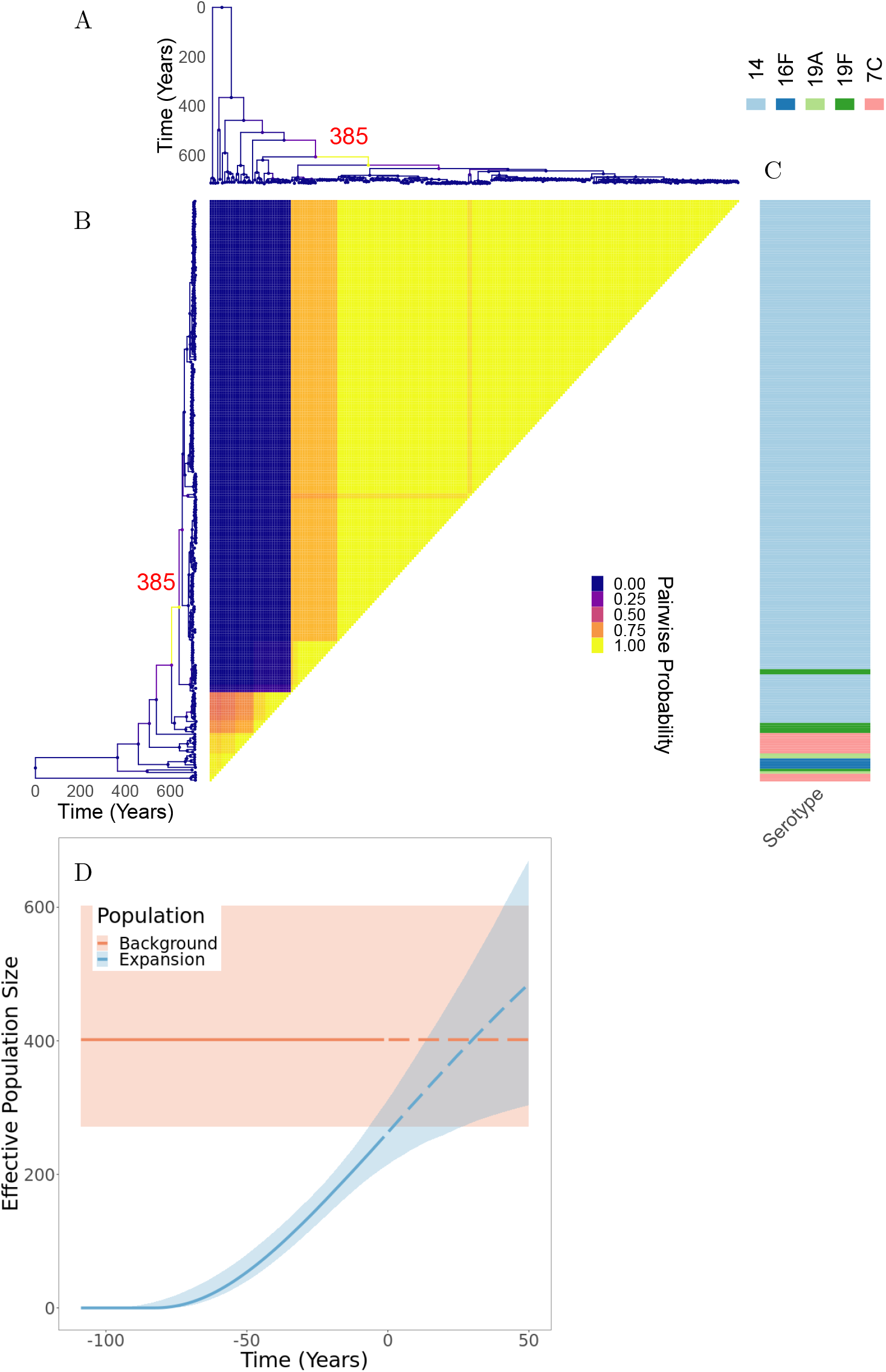
Application to GPSC18 *Streptococcus pneumoniae* phylogeny. (A) Dated phylogeny with branches colored according to the inferred probability of clonal expansion. (B) Pairwise matrix showing the posterior probabilities of any two samples belonging to the same subpopulation. (C) Color map showing serotype values. (D) Posterior summary of the inferred effective population size functions. The colored regions represent 95% credible interval and the lines represent median. Solid denotes past effective population size inference and dashed represents prediction of future effective population size.

### Application to methicillin-resistant *Staphylococcus aureus* dataset

We reanalysed a previously published dataset of genomes of methicillin-resistant *Staphylococcus aureus*(MRSA) from the USA300 lineage (Uhlemann et al. 2014). This lineage was first reported in the early 2000s but quickly spread throughout the United States to become a leading cause of community-acquired skin infections (Challagundla et al. 2018). The dataset consists of 347 genomes isolated between 2006 and 2011, for which we constructed a dated phylogeny using BactDating (Didelot et al. 2018) under the additive relaxed clock model (Didelot et al. 2021). The run time for our clonal expansion analysis software was just under 19 hours for 10^8^ iterations. The results are shown in Figures 6 and S5. The posterior mean for the number of clonal expansions was 3.04, with 28%, 42% and 27% posterior probability assigned to having 2, 3 and 4 clonal expansions, respectively. The most probable posterior population structure therefore consists of three expansions which are nested into one another. The first expansion occurs at branch 374, which then gives rise to an expansion associated with branch 84 and which finally gives rise to expansion starting from branch 217 (Figure 6). The first expansion on branch 374 is the most certain one, and also the most significant one since it splits from the background population which is of a constant population size. This result therefore suggests that it is not the whole of the USA300 MRSA lineage that expanded, but rather a large subset of it which is associated almost perfectly with the presence of the arginine catabolic mobile element (ACME) (Figure 6). ACME provides polyamine resistance as well as other functions (Joshi et al. 2011). An association between ACME and the expansion within USA300 has been suggested before (Uhlemann et al. 2014; Challagundla et al. 2018) but here for the first time we have detected it using a well-suited model of clonal expansion. A previous phylodynamic analysis showed the temporal association between the USA300 growth rate and the consumption of *β*-lactams assumed that the whole population followed the same dynamic function (Volz and Didelot 2018). We show here that this is not correct but this previous analysis remains approximately valid since the vast majority of genomes are part of the ACME-associated clonal expansion. The other two putative expansions that are nested within the first one do not seem associated with a clear genetic change that would provide a selective advantage, but are more likely to correspond to founder effects occurring as USA300 spread in different parts of the human population (Challagundla et al. 2018).

**Figure 6:**
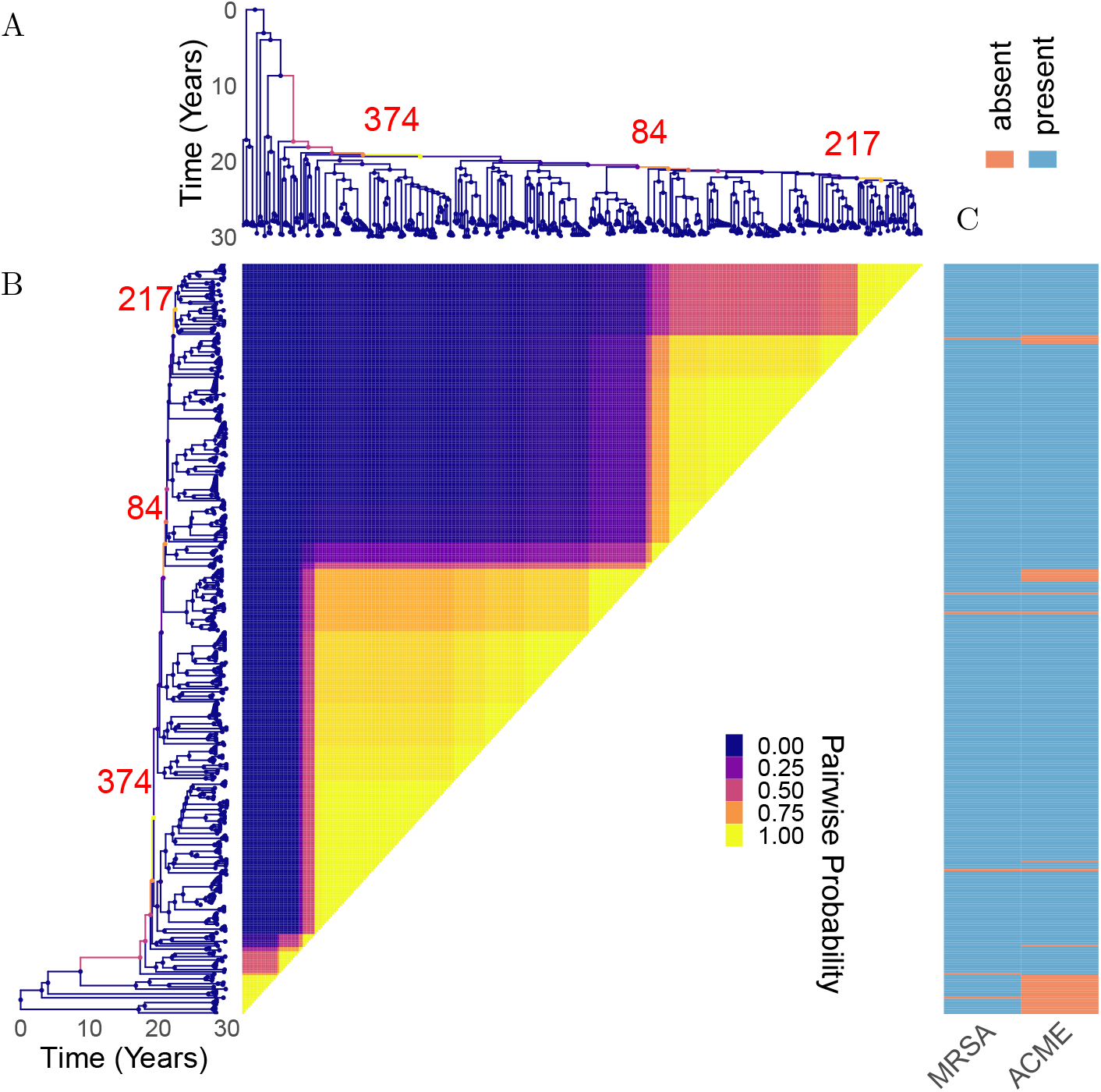
Application to Methicillin Resistant *Staphylococcus aureus* dataset. (A) Dated phylogeny with branches colored according to the inferred probability of clonal expansion. (B) Pairwise matrix showing the posterior probabilities of any two genomes belonging to the same subpopulation. (C) Color map showing the presence of phenotypes associated with virulence.

### Application to *Streptococcus pneumoniae* dataset GPSC9

We also analysed a previously described global collection of genomes from the Global Pneumococcal Sequence Cluster 9 (GPSC9) (Gladstone et al. 2020). This dataset consists of 277 genomes collected between 1995 and 2016 for which a dated phylogeny has been previously published (Gladstone et al. 2020). The MCMC was run for 10^8^ iterations and terminated within 18 hours. The results are shown in Figures 7 and S6. The posterior mean for the number of expansions was approximately 3, with 56% of the posterior probability mass on this number. Approximately 25% of the probability mass rests on a two expansion scenario, and the remainder is distributed between cases with four or more expansions. The latter may be closer to the truth given the previously noted tendency to underestimate clonal expansion numbers (Figure 4). The most certain clonal expansion occurred on branch 389 and corresponds to isolates from all over the world, but are unique within GPSC9 in containing the ermB1 erythromycin resistance gene and being of a serotype not covered by the pneumococcal conjugate vaccines (Figure 7). This clade therefore represents a clear example of vaccine escape by replacement of the capsular locus (Mostowy et al. 2017), followed by worldwide spread. Other identified groups of genomes correspond to locally successful clades as previously described (Gladstone et al. 2020). For example the expansion on branch 288 corresponds to a clade that has successfully established itself throughout the African continent as well as India, with around 50% posterior support to separate the Indian component within this expansion. The background population corresponds to the first South African clade previously identified (Gladstone et al. 2020). These results showcase once again how differences in the phylodynamic trajectories of sublineages are not always caused by a selective advantage of the pathogen, but often linked with the structure of the host population.

**Figure 7:**
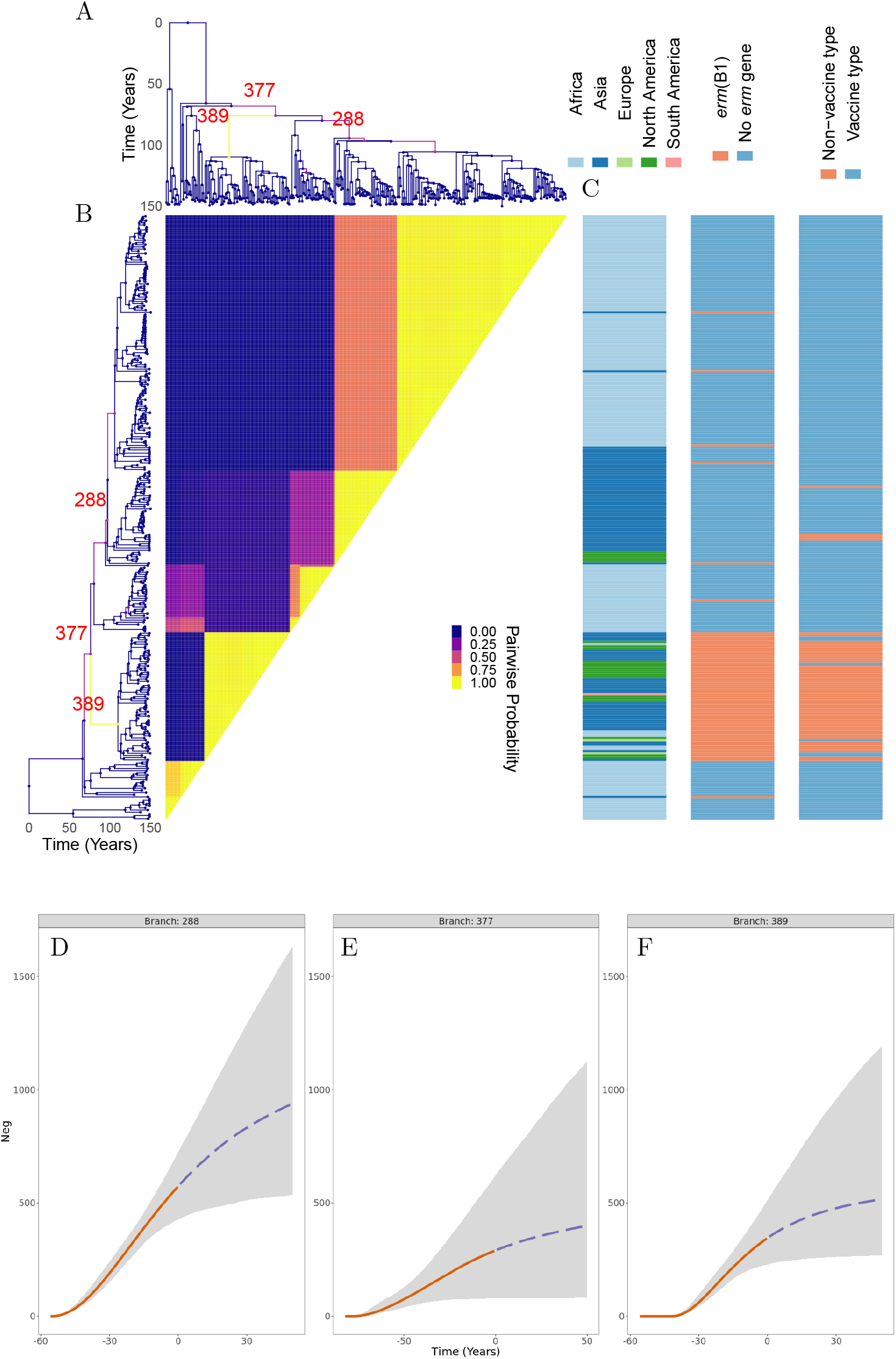
Application to GPSC9 *Streptococcus pneumoniae* phylogeny. (A) Dated phylogeny with branches colored according to the probability of clonal expansion. (B) Pairwise matrix showing the posterior probabilities of any two samples belonging to the same subpopulation. (C) Color map showing geographical sampling location, *erm* gene presence, and whether the serotype is covered by the vaccine. (D-F) Posterior summary of the inferred effective population size functions.

## DISCUSSION

Detecting emerging microbial populations is a persistent and critical public health challenge. However, robust solutions to this problem have been little explored. In this work, we describe a novel, computationally tractable Bayesian approach to finding expanding populations within dated phylogenies. Using simulated phylogenies, we estimated the false positive rate of the approach, which was about 2% in the simulations performed. We also estimated the sensitivity of detection of clonal expansions, which was of the order or 75%, with limited sensitivity attributable to the limited phylogenetic signature left by expansions occurring in antiquity, very recently, or with limited sampling. Importantly, in an analysis of real data from three separate microbial populations causing high burdens of human disease, we identified clonal expansions associated with known virulent factors, drug resistance loci, and absence from vaccine coverage, all biologically credible determinants of clonal expansion. Thus, the application of the approach on both simulated and real world microbial populations indicate the approach described may have wide application. To allow widespread use of our new methodology, we provide an implementation in the form of a R package.

Our methodology has a number of limitations, inherent in the assumptions we have made in our model. Firstly, we assume that the background population, before any clonal expansion occurred, has a constant population size. This assumption would be invalidated for example if the whole population under analysis has been expanding. However, in this case a clonal expansion event would be inferred close to the root. Furthermore, the choice of a constant background population size is convenient from a statistical point of view since it allows scaling of many parameters against the size of the background population (see Table 1). Another choice we made concerns the form of the demographic function after a clonal expansion occurs (Equation 1). Once again this is a choice of convenience, since this function starts at zero when the expansion starts, plateaus at a well-defined carrying capacity value and its reciprocal has an analytical primitive as needed (Equation 6). Our function approximates well the logistic growth behaviour we seek to model and which arises for example in a susceptible-infectious-susceptible SIS model (Allen 2008). Future work could seek to investigate other choices of functions, but choosing another function with similar properties would probably not make much difference to inference results. Our model also assumes that clonal expansions are the only type of phylodynamic events to occur, disallowing for example the possibility for any population size reduction. This is partly because the effect of reduction on phylogenies is less dramatic than sudden growth, so that such events would be harder to detect, but also and mostly because our aim was to provide a method for clonal expansion analysis rather. Further work should seek to expand on our method and develop a more complete framework for the analysis of differential phylodynamic trajectories between lineages, although attention should be given to the identifiability of model parameters.

Biased sampling is a well described confounding factor in phylodynamic studies (Dearlove et al. 2017). To investigate this effect on our method, we simulated standard coalescent trees with many leaves, and then downsampled the leaves in one lineage by a factor that varied between 0.2 and 1. When the bias was strong enough, a clonal expansion event was often detected (Figure S7). However, this behaviour is to be expected, since without any clonal expansion there would be no structure in our model and therefore no explanation for the difference in sampling intensity. Indeed biased sampling can only be achieved if we consider some tree structure, with at least one clade being biased sampled compared to the others. Detecting a clonal expansion event can then be thought of as revealing this underlying structure in the phylogeny, even if in this case there is no underlying difference in the phylodynamic properties between clades.

There are few previous methods to which our approach can be compared, as this is a first-in-class principled approach to the key problem of detecting clonal expansions, whereas the vast majority of existing phylodynamic methods assumes that all lineages follow the same demographic function (Ho and Shapiro 2011). A recent study proposed a non-parametric test of this assumption which can be used to split a phylogeny into separate components but which does not allow further analysis of the phylodynamic properties of each component (Volz et al. 2020). Perhaps the closest existing method is the recently proposed multi-type birth-death (MTBD) model (Barido-Sottani et al. 2020) which is based on the birth-death model (Stadler 2010). In both cases the aim is to model the effect of population heterogeneities in dated phylogenies. However, the model we present is based on a coalescent process as opposed to a birth-death type process, and as such makes fewer assumptions about sampling (Volz and Frost 2014). Furthermore the scenario being modelled is quite different, and is underpinned by a completely different set of assumptions. Since our focus is specifically on clonal expansions, an equivalent to birth-death changes only occurs when all members of a given clonal expansion have coalesced, which is not the case with the MTBD model (Barido-Sottani et al. 2020). Instead, our model is more closely related to the multi-species coalescent (Degnan and Rosenberg 2009), but with the key differences that we consider the phylogeny of just a single locus, and that there is an extreme bottleneck at speciation events. Some comparison may also be drawn with genetic clustering based on fitting a Markov-modulated Poisson process (MMPP) (McCloskey and Poon 2017), although this method focuses on detecting small scale outbreaks, whereas we are interested in a phylodynamic behaviour on a significantly larger scale. Furthermore, the assumptions are completely different: our model is phylodynamic and does not represent an approximation of a transmission tree. Finally, our method is related with approaches to detecting structure which are not based only on the phylogeny, but exploit integration with other type of data (Baele et al. 2016), for example using the distribution of a phenotype (Ansari and Didelot 2016) or the geographical origin of the samples (Bloomquist et al. 2010).

The approach presented here should be applicable to a wide range of microbes, as long as their ancestral process can be summarised using a dated phylogeny, and that the genomic data is sufficiently informative to reconstruct such a tree with reasonable accuracy. Our method was designed primarily to analyse retrospectively the structure of microbial populations, as illustrated in the three applications to real life datasets we described. However, our method could also be useful in a public health setting to detect, confirm and analyse suspected outbreaks of infectious diseases, or the emergence of new lineages with increased transmissibility, bearing in mind that clonal expansion events can also be associated with non-epidemic factors.

## ACKNOWLEDGEMENTS

This work was supported by the UK Engineering and Physical Sciences Research Council (EPSRC) grant EP/S022244/1 for the EPSRC Centre for Doctoral Training in Mathematics for Real-World Systems II. We acknowledge funding from the National Institute for Health Research (NIHR) Health Protection Research Unit in Genomics and Enabling Data.

