## Supplementary Material for "Bayesian inference of clonal expansions in a dated phylogeny"

### Bayesian inference of clonal expansions in a dated phylogeny Supplementary Material

#### Updates of the reversible jump MCMC

For inference we use reversible jump MCMC in order to explore the space of models with varying number of clonal expansions. The MCMC moves consist of five within model moves, as well as one transdimensional move that adds or removes clonal expansions.
We will be using the structure of the tree to guide our proposal distributions. It is worth noting that divergence of a clonal expansion effectively occurs along a branch. We can justify this by noting that a given tip partition corresponding to a certain colour must fully coalesce with itself up to the MRCA of the expansions before changing colour to that of its parent. This is because the effective population size vanishes as the divergence time reached. The branch along which the expansion diverges is the one leading into the expansion MRCA. It remains to demonstrate that the the divergence time for a expansion cannot exceed the time of parent of this branch. This is straightforward as the divergence time being greater than the parent of the branch leading into the expansion MRCA would imply that partitions with different colouring coalesced - an event that happens with probability zero.
In order to efficiently explore all tip partitions compatible with the genealogy given, tip partitions are generated by selecting a branch along which an emergence of given colour is supposed to happen. All tips corresponding to the clade that has the child node along the branch as its MRCA are then assigned to the given colour, with the exception of tips that correspond to clades that diverge earlier.

##### Within Model Moves

We will use the following notation to describe updates.
Let

$$x^{(M)}_{r_j \to r'} \tag{S1}$$

Denote the update of rate parameter $r_j$ corresponding to the $j$-th clonal expansions to the value $r^{'}$ in the $M$-expansion model. Analogously, all within model updates will be denoted this way.

The within model updates will now be described.
The first move is:

$$x^{(M)}_{(h_j, N_j) \to \left(h', N'\right)} \quad \forall j < M \tag{S2}$$

This move updates simultaneously the half-life and the carrying capacity of the expansion. The new values are distributed according to

$$\left(h^{'}, N^{'}\right) \sim (\mathcal{N}(h_j, \sigma_r), \texttt{lognorm}(log(N_j), \sigma_K)) \tag{S3}$$

The proposal ratio of this move is:

$$\frac{f_{ln}(N_j; log(N^{'}), \sigma_K)}{f_{ln}(N'; log(N_j), \sigma_K)} \tag{S4}$$

Where $f_{ln}(x; a, b)$ stands for the value of the lognormal distribution pdf with log-mean $a$ and log-standard deviation $b$ evaluated at $x$.

The second move is:

$$x^{(M)}_{(\mathbf{f}, t_{div_j}, D) \rightarrow (\mathbf{f}', t'_{div}, D')} \tag{S5}$$

This moves updates the colouring of the leaves and thus the memberships of clonal expansions, as well as the time of the expansion. This is perhaps the most complicated move and proceeds as follows:
Let $b_j$ denote the branch along which the divergence event for the $j$-th expansion lies, i.e. the branch leading into the MRCA of the expansion, and with $m_j$ the MRCA of the expansion. Let $t_\Delta$ be a normally distributed variable centred at $t_{div}$, $t_\Delta \sim \mathcal{N}(t_{div}, \sigma_t)$. Define $dir = 1$ if $t_\Delta > t_{div}$ and $dir = -1$ otherwise. If $t_\Delta$ is greater than the time of the root $t_{root}$, set $t^{'}_{div} = t_{root} - (t_\Delta - t_{root})$ so that the root acts as a reflecting boundary. Otherwise set $t^{'}_{div} = t_\Delta$. Set $b^{'} = b_j$, $u = 0$, $v = 0$. The following process gets repeated until $t^{'}_{div}$ lies within $b^{'}$, i.e. is less than the time of its parent node and greater than the time of its child.

If $dir > 0$ we move towards the root of the tree as follows. If the parent of $b^{'}$ is the root, set $b^{'}$ to be the other child branch exiting the root and $dir = -1$. Otherwise, Set $b^{'}$ to be the edge that terminates in the parent of $b^{'}$ and set $u = u + 1$.

If $dir < 0$ we move away from the root of the tree. With equal probability 1/2 set $b^{'}$ to be one of the two branches exiting the child node of $b^{'}$ and set $v = v + 1$.

Finally the new branch as $b^{'}$, and the new MRCA, that is the child node along branch $b^{'}$, as $m^{'}$. Colour the tips corresponding to the clade defined by $m^{'}$, with the exception of tips corresponding to clades that diverge earlier as described previously in this section. Finally, set the parent populations $D^{'}$ so that they are compatible with the updated divergence times and tip partition. This is done for each expansion $i$, by setting $d_i$ to be equal to the lineage membership of the parent node of the MRCA of expansion $i$.
The proposal ratio for this move is:

$$\frac{2^v}{2^u} \tag{S6}$$

The third move updates the probabilities vector $\boldsymbol{\theta}$:

$$x^{(M)}_{\boldsymbol{\theta} \rightarrow \boldsymbol{\theta}'} \tag{S7}$$

For this move first simulate indices $k, l \sim U(\{1...M\})$. Next simulate $u \sim U((0, \theta_k))$. Set $\theta^{'}_i = \theta_i \quad \forall i \in \{1...M\} \setminus \{k, l\}$ and $\theta^{'}_k = \theta_k - u$, $\theta^{'}_l = \theta_l + u$.
The proposal ration for this move is:

$$\frac{\theta_k}{\theta^{'}_l} \tag{S8}$$

The fourth and final move updates the effective population size $N_M$ of the parent population:

$$x^{(M)}_{N_M \rightarrow N'} \tag{S9}$$

With distributed according to

$$N^{'} \sim \mathcal{N}(N_M, \sigma_N) \tag{S10}$$

The proposal ration for this move is 1.

#### Transdimensional moves

The move that removes an expansion $x^{(M)\to(M-1)}$ proceeds by selecting an expansion index $k$ uniformly at random from $\{1...M-1\}$ ($M-1$ as parent population cannot be removed), removing the associated parameters from the parameter vector and setting $M^{'} = M-1$. The tip partition $\mathbf{f}$ is updated by colouring tips according to the scheme mentioned in the previous section using the new parameter vector with the $k$-th expansion parameters removed. The probability vector $\boldsymbol{\theta}^{'}$ is updated by first selecting $l$ uniformly at random from $\{1...M^{'}\}$, and setting $\theta_i^{'} = \theta_i \quad \forall i : i < k, i \neq l$, $\theta_i^{'} = \theta_{i+1} \quad \forall i : i \geq k, i \neq l$ and finally $\theta_l^{'} = \begin{cases} \theta_l + \theta_k & l < k \\ \theta_{l+1} + \theta_k & l \geq k \end{cases}$.
The proposal ratio for this move is:

$$\frac{(M-1) \times (M-1) \times \pi(N^{'} \mid N_M) \times \pi(t_{div}^{'} \mid N_M) \times \pi(r^{'})}{\theta_l^{'} \times (M-1)} \tag{S11}$$

The reverse move that adds an expansion $x^{(M)\to(M+1)}$ proceeds by setting $M^{'} = M+1$ and simulating expansion parameters from the priors as:

$$\begin{aligned} N^{'} &\sim \pi(N \mid N_M) \\ t_{div}^{'} &\sim \pi(t_{div} \mid N_M) \\ r^{'} &\sim \pi(r) \end{aligned} \tag{S12}$$

The new expansion is added under the index $M^{'} - 1$. The tip partition $\mathbf{f}$ is updated by adding a new colouring, such that it is not identical to an existing colouring, and so that the clade defined by this colouring coalesces with itself before $t_{div'}$. Furthermore the parent clade $d^{'}$ must be selected so that the MRCA of the diverging clade coalesces with a node of the parent clade after $t_{div'}$. This can be done by first selecting a branch $b'$ uniformly at random from all branches with child node before $t_{div'}$ and parent node after $t_{div'}$. Next, setting the new partition to be all leaves belonging to the clade with the MRCA given by the child node of $b'$ excluding those that belong to diverging clades with time of divergence before $t_{div'}$. Finally, the parent population vector $D^{'}$ has to be updates by for expansion $i$ setting $d_i^{'}$ to be equal to the expansion membership of the parent node of the MRCA of expansion $i$. The probability vector $\boldsymbol{\theta}^{'}$ is updates by selecting $k$ uniformly at random from $\{1...M\}$, and $u \sim U((0, \theta_k))$. Set

$$\theta_i^{'} = \begin{cases} \theta_i & i < M, i \neq k \\ \theta_i - u & i < M, i = k \\ u & i = M \end{cases} \tag{S13}$$

Then set

$$\theta_{M'}^{'} = \begin{cases} \theta_M & k \neq M \\ \theta_M - u & k = M \end{cases} \tag{S14}$$

The proposal ration for this move is:

$$\frac{\theta_k \times (M)}{(M) \times (M) \times \pi(N' \mid N_M) \times \pi(t'_{div} \mid N_M) \times \pi(r')} \tag{S15}$$

When selecting a move, we first select whether to do a within model move or a transdimensional move with probability 1/2. Next if within model moves are selected, we undertake one of the within model moves with equal probability. If transdimensional moves are selected, we choose whether to increase or decrease the number of expansions with probability 1/2.

### Prior Rationale

The starting point for the choice of priors is using the background population as scale parameter for the entire process. For the background population size, a relatively non-informative log-normal distribution is used. For the number of expansions, we use a Poisson distribution as a prior. This allows for a simple parametrisation, and is consistent with the assumption that clonal expansions are relatively rare. We use a Dirichlet distribution with a single concentration parameter as a prior distribution on membership probabilities. By setting this parameter to be greater than one, we introduce slight bias against population assignments that are exceedingly unbalanced and only assign a very small proportion of tips to an expansion. A Gamma distribution is used to impose a prior on the time of clonal expansions. The parametrisation used is a closely related to the method of moments parametrisation, with the key difference that the moments are expressed in terms of a proportion of the background population size, as this parameter is in general tied to the entire temporal scale of the process. Typical values of $\kappa, \nu$ should be chosen such that the distribution remains dispersed and non-informative, while also placing more density closer to the more recent sampling times rather than the expected time of the tree root assuming background population being constant. This is to reflect the number of extant lineages decreasing with time. Finally, an exponential prior is imposed on the time it takes for an expansion to reach half its carrying capacity. This is parametrised so that the mean is equal to fraction $1/\lambda_r$ of the background population size. We chose an exponential distribution assign lower likelihood to more rapid population growth as this is generally likely to be less feasible biologically.

### Biased Sampling Simulation

We first sample 40 coalescent trees under constant population size, each with 500 tips. Within each tree, a particular clade is chosen so that the MRCA of this clade occurs roughly at half the height of the tree. Tips are then removed from this clade with probability $p$, and with probability 0.5 outside of the the clade. The clade therefore suffers from a downsampling bias of $2p$ relative to the rest of the tree.

A

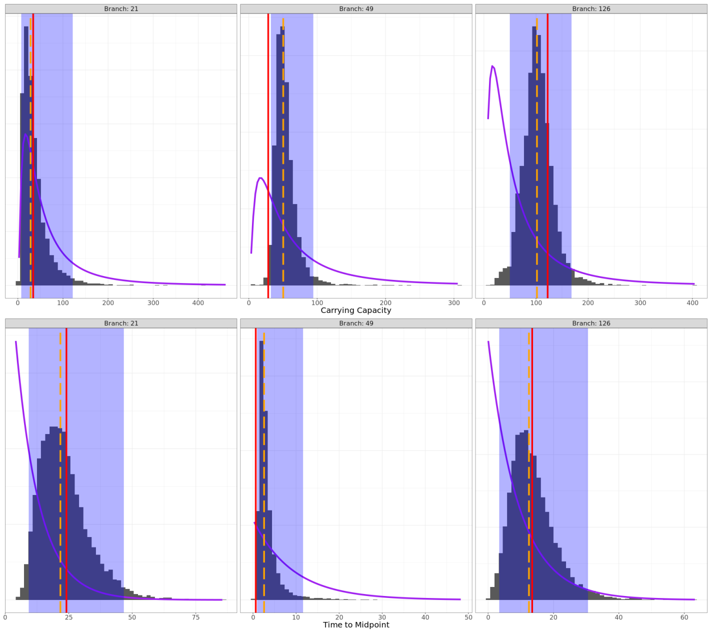


B

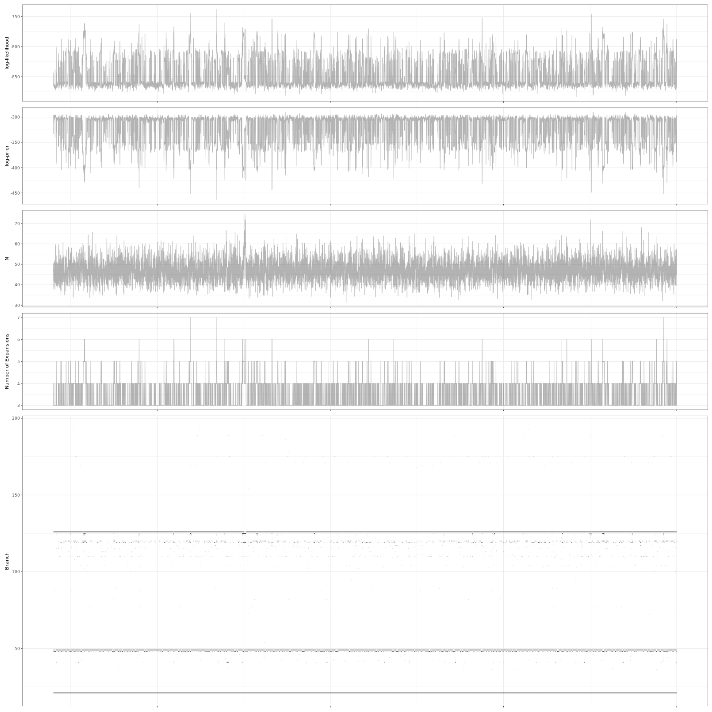

Figure S1: Supplementary information for the analysis of the first simulated dataset. (A) Parameter marginals for the three highlighted expansions. Expansion branch in header. True value denoted by solid red bar. Median denoted by dashed orange line. 95% Credible interval shaded in blue. Purple line represents prior density. (B) Traces for likelihood values, prior values, background population size, number of expansions, and expansion position.

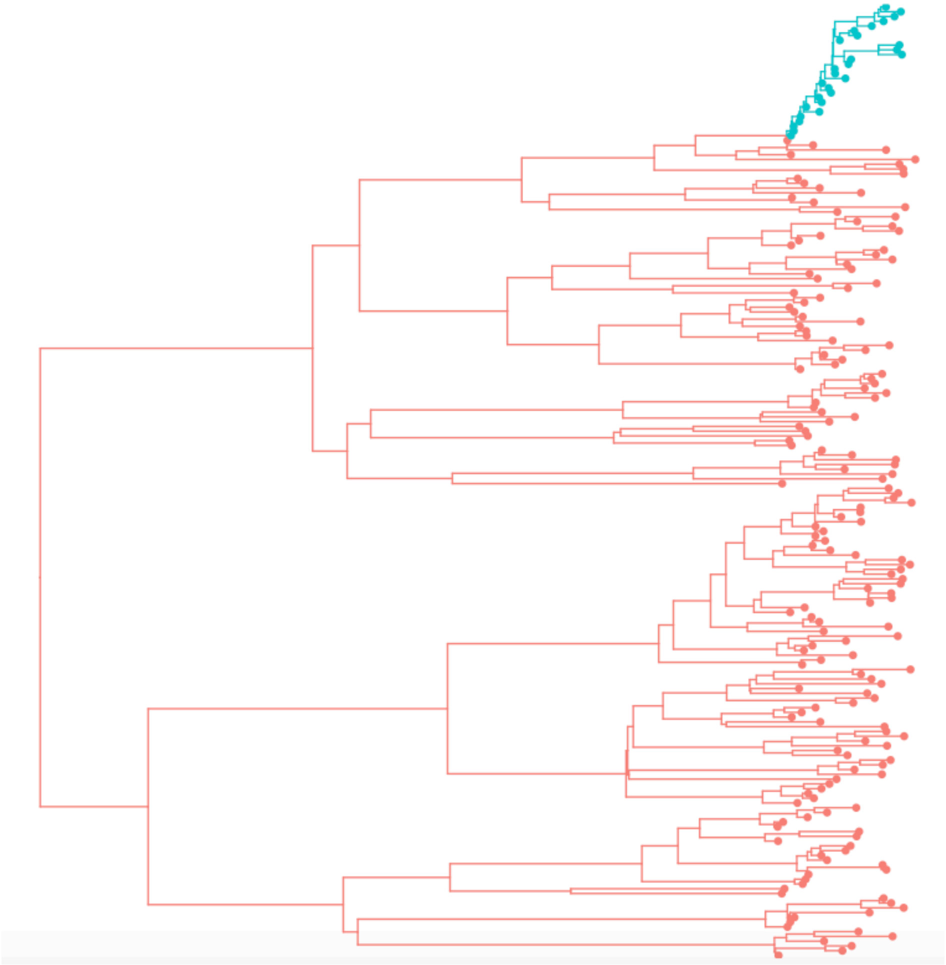

Figure S2: Application of treestructure to the first simulated dataset.

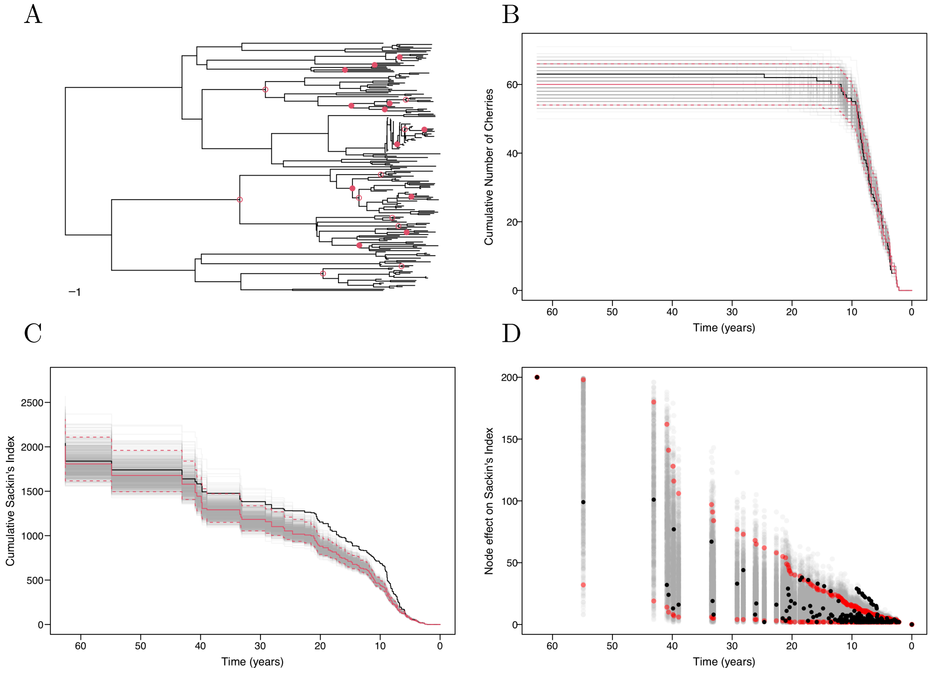


Figure S3: Application of treeImbalance to the first simulated dataset. (A) Dated tree with nodes identified as significantly more asymmetric than expected with the Bonferroni correction are marked with a filled red circle and those which are significant at an unadjusted p-value of 5% are shown by an open red circle. (B) Observed cumulative number of cherries over time (black), with results from permuted trees (grey). Red lines show the medoid (solid) and 95% confidence interval of the permuted results (dashed). (C) Trajectories for the cumulative Sackin's index. (D) Node effect on Sackin's index over time.

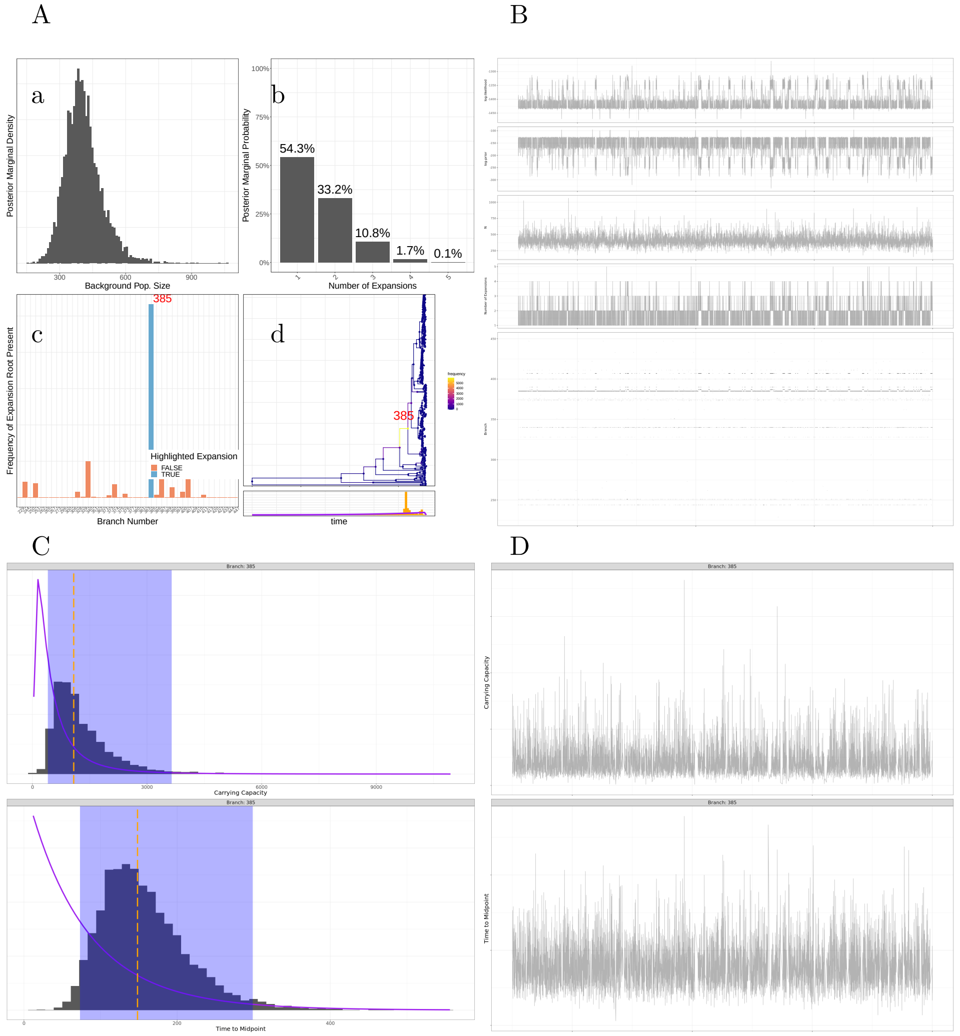


Figure S4: Supplementary information for the analysis of the GPSC18 dataset. (A) - Summary plot displaying: (a) Background population size marginal histogram. (b) Number of expansions marginal histogram. (c) Marginal histogram of probabilities that an expansion lies along a given branch. (d) Date phylogeny with branches coloured according to the probability that an expansion lies along it. (B) - Traces for likelihood values, prior values, background population size, number of expansions, and expansion position. (C) - Parameter marginals for branches with the highest posterior probabilities of expansion, as marked by red numbers in (A). Median denoted by dashed orange line. 95% Credible interval shaded in blue. Purple line represents prior density. (D) - Parameter traces for branches with the highest posterior probabilities of expansion, as marked by red numbers in (A).

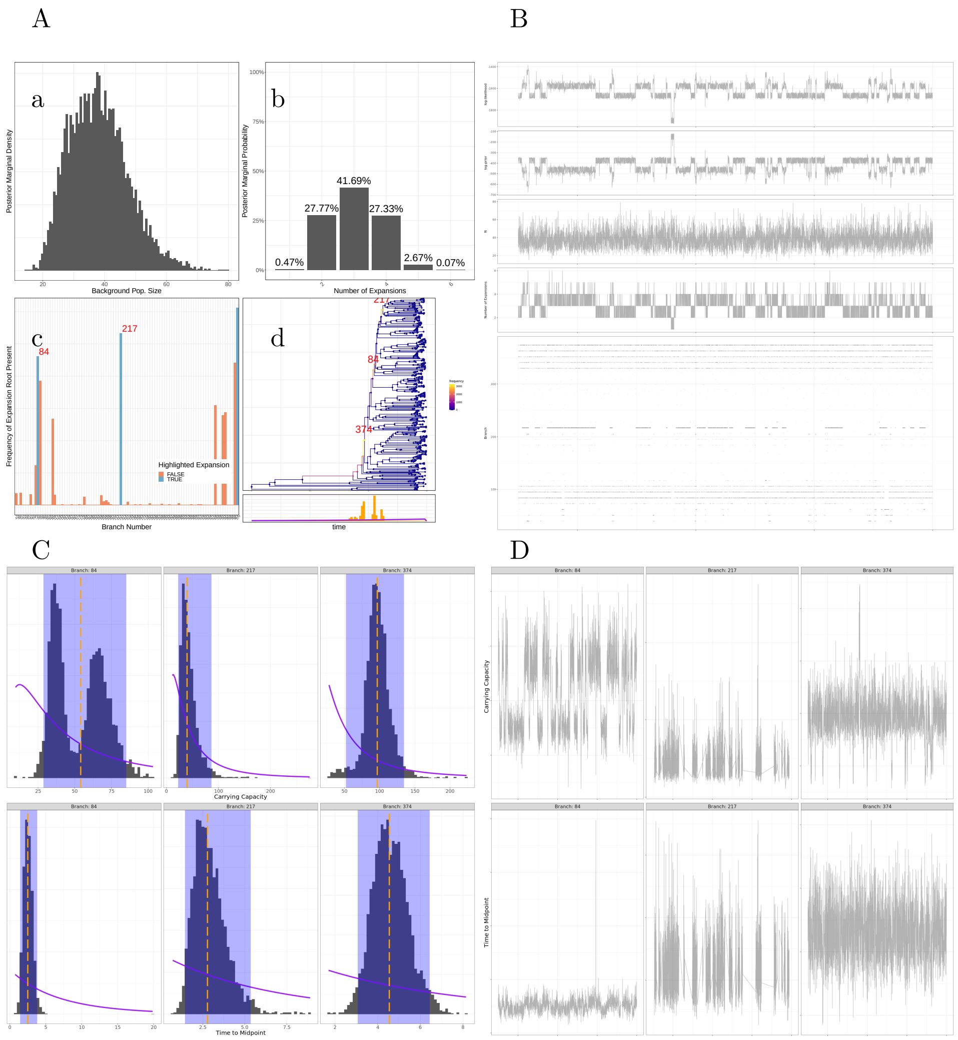


Figure S5: Supplementary information for the analysis of the MRSA dataset. (A) - Summary plot displaying: (a) Background population size marginal histogram. (b) Number of expansions marginal histogram. (c) Marginal histogram of probabilities that an expansion lies along a given branch. (d) Date phylogeny with branches coloured according to the probability that an expansion lies along it. (B) - Traces for likelihood values, prior values, background population size, number of expansions, and expansion position. (C) - Parameter marginals for branches with the highest posterior probabilities of expansion, as marked by red numbers in (A). Median denoted by dashed orange line. 95% Credible interval shaded in blue. Purple line represents prior density. (D) - Parameter traces for branches with the highest posterior probabilities of expansion, as marked by red numbers in (A).

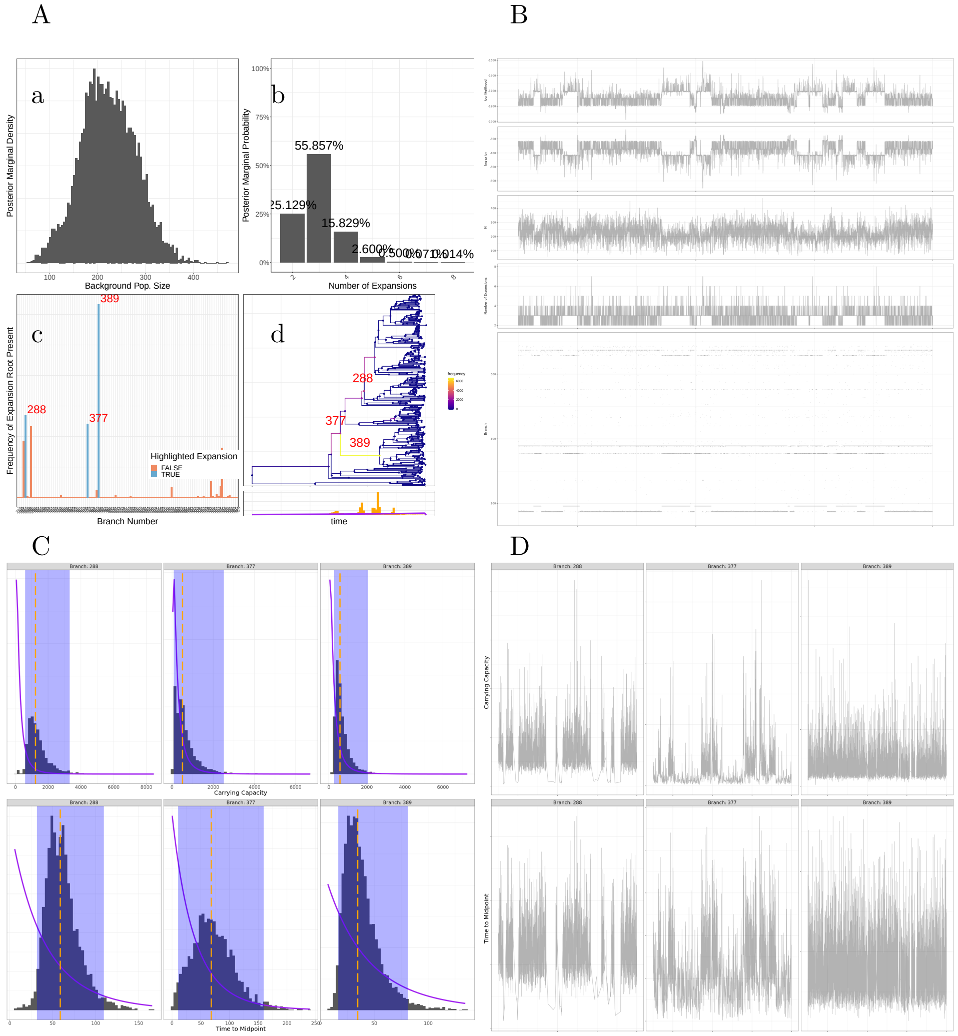


Figure S6: Supplementary information for the analysis of the GPSC9 dataset. (A) - Summary plot displaying: (a) Background population size marginal histogram. (b) Number of expansions marginal histogram. (c) Marginal histogram of probabilities that an expansion lies along a given branch. (d) Date phylogeny with branches coloured according to the probability that an expansion lies along it. (B) - Traces for likelihood values, prior values, background population size, number of expansions, and expansion position. (C) - Parameter marginals for branches with the highest posterior probabilities of expansion, as marked by red numbers in (A). Median denoted by dashed orange line. 95% Credible interval shaded in blue. Purple line represents prior density. (D) - Parameter traces for branches with the highest posterior probabilities of expansion, as marked by red numbers in (A).

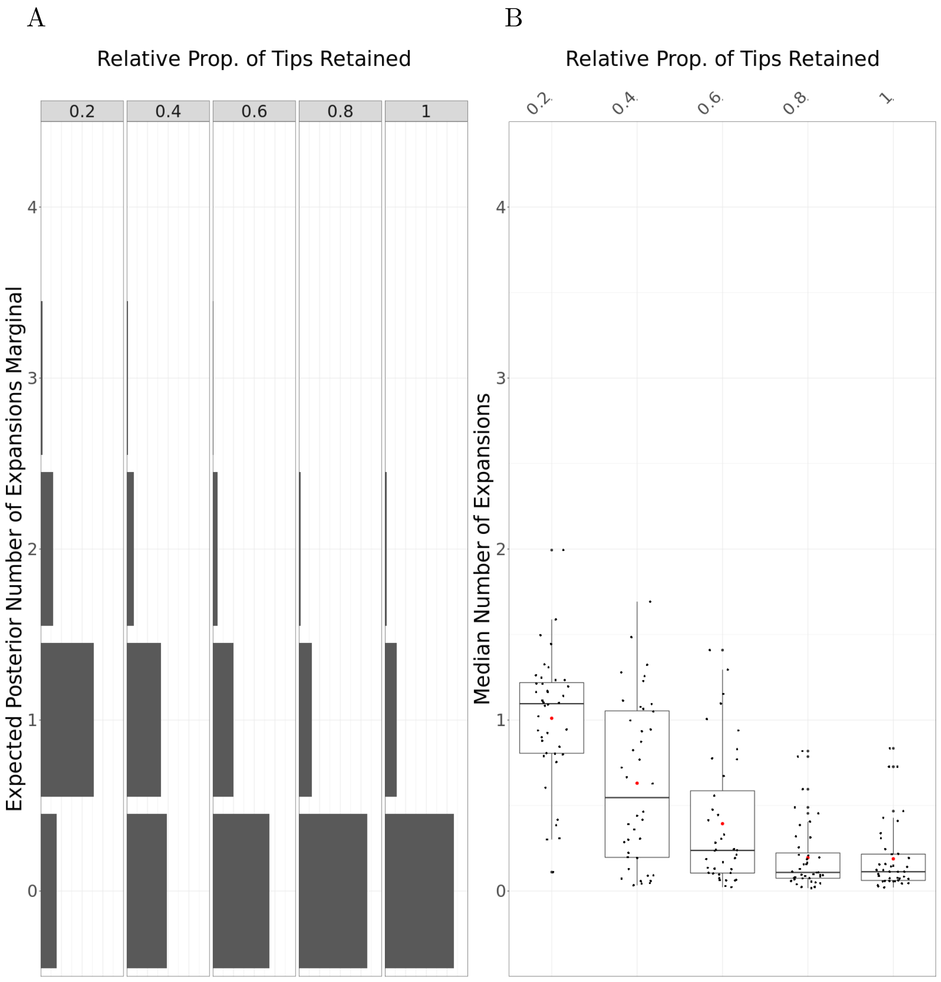


Figure S7: Supplementary information for biased sampling simulation. (A) - Panel displaying expected number of expansions marginal distribution, for different levels of biased sampling. (B) - Box plot of the posterior medians of the number of expansions.
